# The divergent effects of CDPPB and cannabidiol on fear extinction and anxiety in a predator scent stress model of PTSD in rats

**DOI:** 10.1101/542118

**Authors:** John Shallcross, Peter Hámor, Allison R. Bechard, Madison Romano, Lori Knackstedt, Marek Schwendt

**Author notes:** Correspondence: Marek Schwendt.

## Abstract

Post-traumatic stress disorder (PTSD) is a disorder with no clear FDA-approved treatments that reduce symptoms in the majority of patients. PTSD individuals possess an impaired capacity for extinguishing fear memory associations. As such, considerable focus has been given to the development of extinction-enhancing pharmacological agents to be used in combination with PTSD treatments. Here we use a predator-threat animal model of PTSD to test the ability of two compounds to enhance contextual fear extinction and reduce anxiety. Mirroring the heterogeneity observed in human response to trauma, an exposure to predator threat, including the fox odor TMT, have been shown to induce long-term changes in anxiety behavior in only subsets of “susceptible” rats. Here, two weeks following a ten-minute exposure to a predator odor, rats were classified into stress-Susceptible (Sus) and stress-Resilient (Res) phenotypes using cut-off behavioral criteria for elevated plus maze and acoustic startle response performance. One week following classification, Sus rats underwent three days of context fear extinction. We found that Sus rats increased freezing from day one to day two. Treatment with the mGlu5 positive allosteric modulator CDPPB, but not the phytocannabinoid cannabidiol (CBD), prior to sessions resulted in reduced freezing. CDPPB administration resulted in an increase of Fos immunoreactive cells in the medial prefrontal cortex, indicative of increased neuronal activity. Finally, we used the light-dark box test to measure phenotypic differences and the effects of CDPPB and CBD on unconditioned anxiety two weeks after classification. We found that Res rats showed less anxiety compared to Sus rats, and that CBD, but not CDPPB, administered prior to testing was anxiolytic in Sus rats. Taken together, the present data indicate that mGlu5 PAMs such as CDPPB hold promise for treating human PTSD patients as they enhance extinction of fear without increasing general anxiety. Polytherapy with medications such as CBD may be necessary in order to attenuate general anxiety and future directions will explore this hypothesis.

## Introduction

Post-traumatic stress disorder (PTSD) develops in a subset of individuals following a traumatic event (Perkonigg et al., 2001). A characteristic feature of PTSD is impaired fear memory extinction (Guthrie & Bryant, 2006; Orr et al., 2000), which contributes to the persistent anxiety and hyperarousal experienced by affected individuals (Norrholm et al., 2011; Herman, 1992). Fear extinction is an active learning process where stimuli that previously elicited fear are repeatedly presented in the absence of threat and gradual reduction fear responses occurs (Bouton et al., 2006). While extinction-based exposure therapies are frequently used as a strategy for treating anxiety-like disorders, PTSD associated extinction deficits compromise the efficacy and outcomes of these treatments (Schottenbauer et al., 2008). Consequently, there is an urgent need to improve currently available therapies for PTSD. One approach that directly addresses extinction deficits in PTSD advocates for the adoption of extinction-enhancing pharmacological agents that can be co-administered with exposure therapy to improve treatment outcomes (e.g. Rothbaum et al., 2014).

In this regard, animal models are essential to study the neurobiology underlying fear extinction and for the development of novel extinction-enhancing therapeutics. The most commonly used models are grounded in Pavlovian fear conditioning principles (Pavlov, 1927; Rescorla, 1988). Fear conditioning involves pairing an unconditioned aversive stimulus (US; e.g. mild electric shock) with neutral conditioned stimuli (CS; e.g. a discrete cue or context) until a conditioned fear response (CR; e.g. freezing, changes in heart rate) is produced following delivery of the CS alone. Like exposure therapy, fear extinction training involves prolonged, or repeated presentations of the CS alone, and ideally results in the gradual elimination of the CR (Barad, 2005; Rothbaum & Schwartz, 2002).

Most animal models used to investigate the neural mechanisms of fear extinction utilize footshock stressors. However, for the study of extinction in the context of PTSD, these models lack face validity, as footshock is not ethologically relevant to rodents and does not mimic induction of PTSD in humans. In addition, the incidence of PTSD diagnoses in only a subset of trauma exposed individuals is not considered in these models, as the consequences of treatment manipulations are assessed in the entire population of conditioned animals and individual differences in anxiety are not accounted for. Importantly, while animals extinguish conditioned fear responses to footshock-conditioned cues, extinction in humans with PTSD is not generally achieved (VanElzakker et al., 2013). Alternatively, exposure to species-relevant predator threat provides an ethologically relevant life-threatening experience that can elicit robust fear responses (e.g. freezing; Wallace & Rosen, 2000), and supports contextual fear conditioning (Endres & Fendt, 2007). Predator threat exposure has also been demonstrated to more closely resemble a PTSD-like profile in that the persistent manifestations of anxiety and hyperarousal following exposure to threat are observed only in a subset of “Susceptible” exposed rats and not in “Resilient rats” (Cohen & Zohar, 2004; Schwendt et al., 2018). Despite these advantages, little is understood about the neural mechanisms involved in the extinction of innate predator fears, or the benefits of extinction-enhancing agents under these conditions. We have previously demonstrated that a single 10-minute exposure to the fox odor 2, 3, 5-Trimethyl-3-thiazoline (TMT) gives rise to distinct stress-Susceptible and Resilient rat phenotypes, each with distinct behavioral, hormonal, and molecular signatures (Schwendt et al., 2018). Notably, we found that only stress “Susceptible” rats displayed increased freezing upon re-exposure to the stress/trauma context whereas “Resilient” rats did not, indicating that inter-subject phenotypic heterogeneity influences the observed outcomes across entire populations of stressed animals. Furthermore, we also demonstrated that Susceptible rats <u>do not</u> decrease freezing over the course of 5 days of extinction to the stress context (Schwendt et al., 2018). Taken together, this data indicates that knowledge gained from classic footshock conditioning experiments may not apply to models that consider differential stress vulnerability.

In healthy humans, neuroimaging studies have revealed an important role for activity in the medial prefrontal cortex (mPFC) during extinction indicated by a rise in the ventral medial prefrontal cortex (vmPFC) coupled with reduced activity in the dorsal lateral prefrontal cortex (dlPFC; Milad et al., 2007) and amygdala (Labar et al., 1998). Opposite patterns are demonstrated in humans with PTSD, with low vmPFC activity and high activity in both dlPFC and amygdala (Milad et al., 2009). As noted above, the neural correlates of fear extinction in rodents have been extensively studied using footshock models, and suggest a conserved mechanism also involving functional interactions between the mPFC and amygdala. In the rodent mPFC, the prelimbic (PL) and infralimbic (IL) cortices (analogous to the dlPFC and vmPFC in humans, respectively) are strongly interconnected with the basolateral amygdala (BLA; Hoover & Vertes, 2007). The BLA is required for extinction of conditioned footshock (Falls et al., 1992), and serves to regulate fear response through output to the central amygdala (CeA) and brainstem regions (Royer et al., 1999; Haubensak et al., 2010). Chemogenetic, or electrical stimulation of IL or PL pathways targeting the BLA reveal opposing influences (Herry et al., 2008; Senn et al., 2014), with IL enhancing, and PL impairing extinction (Sierra-Mercado et al., 2011). Additionally, inhibitory and excitatory IL and PL projections (respectively) regulate BLA excitability, stabilizing fear response inhibition (Cho et al., 2013). This evidence suggests that extinction of footshock conditioned fear requires a switch from PL to IL mediated reciprocal signaling through the BLA. Indeed, the assessment of neuronal activity using c-Fos immunoreactivity reveals high Fos expression in the IL, but not PL following extinction, and PL and BLA Fos expression correlating with extinction resistance (Knapska & Maren, 2009). Few studies have examined fear extinction associated neural function using predator-threat models. Notably, BLA inactivation reduces fear response to a cat or cat odor conditioned context (Takahashi et al., 2007; Bindi et al., 2018), and compared to sham exposed rats, enhanced ΔFosB expression was observed in the mPFC but not BLA of cat odor exposed rats following five-day context extinction (Mackenzie et al., 2010).

Given that fear extinction involves neuronal plasticity, modulation of glutamate receptors has been the focus of many efforts to develop extinction-enhancing agents. mGlu5 receptors regulate bidirectional synaptic plasticity in fear associated brain regions such as the mPFC and BLA (Niswender & Conn, 2010). Pharmacological and genetic inhibition of mGlu5 function impairs extinction of both cues and contexts paired with footshock in rodents (Sethna & Wang, 2016; Fontanez-Nuin, 2010; Sepulveda-Orengo et al., 2013; Xu et al., 2009), while administration of of mGlu5 positive allosteric modulators (PAM) enhances extinction of footshock-associated contexts (e.g. Sethna & Wang, 2014). While mGlu5 antagonists inhibit fear extinction, they also produce anxiolytic effects (Rahman et al., 2017; Porter et al., 2005). The effects of glutamate receptor modulating drugs on extinction to innate predator threat remains largely unknown. One recent study demonstrated that partial agonism of NMDA receptors with D-cycloserine enhanced extinction to a cat odor exposed context (Saridoğan, 2015). Notably, we recently found differences in mGlu5 mRNA expression between stress Susceptible and Resilient rats. As described above, in this study, we separated stress-Susceptible from Resilient rat phenotypes one week after a single TMT exposure based on performance across multiple anxiety measures. Three weeks after the initial TMT exposure, we observed increased expression of mGlu5 and cannabinoid receptor mRNA (CB1) mRNA in the amygdala, suggesting the contribution of these signaling systems in avoiding persistent, trauma induced anxiety. We also reported that in a separate cohort of Susceptible rats that underwent cocaine self-administration, daily treatment with CDPPB did not facilitate extinction of the TMT context, but instead augmented fear (freezing). The effects of CDPPB on fear extinction in cocaine-naïve rats in this model remain undetermined and hence was the primary goal of the present work.

Like mGlu5, CB1 receptors are abundantly expressed in the BLA and mPFC and are important modulators of fear and anxiety signaling (Chhatwal & Ressler, 2007). Human studies have revealed dysregulated expression of CB1 and endogenous CB1 ligands in subjects with PTSD (Neumeister et al., 2013; Hill et al., 2013). In rodents, genetic or pharmacological inhibition of CB1 impairs extinction (Marsicano et al., 2002), while CB1 agonists enhance extinction (Chhatwal et al., 2005; Campolongo et al., 2009). However, CB1 agonists can produce biphasic anxiogenic and anxiolytic effects (Haller et al., 2004; Sink et al., 2010), which may compromise their clinical usefulness. Several recent studies have demonstrated the low-affinity CB1 indirect agonist cannabidiol (CBD), a component of cannabis which lacks THC-like psychoactive effects (Campos et al., 2012a), as a potential PTSD treatment. Notably, CBD can enhance extinction and reduces post-trauma anxiety in both humans and rodents (Das et al., 2013; Bitencourt et al., 2008).

Here, we examined the effect of CDPPB and CBD on the extinction of predator odor (TMT)-induced contextual fear in stress Susceptible rats. We hypothesized that potentiation of mGlu5 or CB1 receptor activity would facilitate extinction. Interestingly, we found that only CDPPB produced extinction enhancements. To explore the neurobiology underlying this effect, brain tissue was collected after the third extinction session and expression of Fos was analyzed in PL, IL, and BLA regions. Given the involvement of PL, IL, and BLA neuronal activity in extinction to conditioned footshock (Cho et al., 2013), and evidence indicating an important role for mGlu5 (Sethna & Wang, 2016), we hypothesized that treatment with CDPPB would result in greater Fos in all three regions. We also expected that due to the opposing influences of the PL and IL neuronal activity during extinction (Vidal-Gonzalez et al., 2006), Fos expression across mPFC regions would be predictive of fear extinction performance. Last, we examined the effects CDPPB and CBD on anxiety in the light-dark box test. Previous studies have found that CDPPB produces either no effect or a moderate anxiogenic effect in previously unstressed rodents (Sethna & Wang, 2014; Lee et al., 2018). As CDPPB enhances extinction of fear, and drugs in this class of mGlu5 PAMs are viable treatments for the enhancement of exposure therapy, it is essential to characterize its effects in Susceptible populations prior to determining whether CDPPB would be a potential treatment for PTSD patients. We predicted that CDPPB treatment would either produce no effect or modestly increase anxiogenic behavior in the light-dark box. However, we hypothesized that CBD treatment would produce anxiolytic effects in the light-dark box, in agreement with previous work demonstrating CBD mediated anxiety reductions in the light-dark box test in rats with a prior exposure to footshock stress (Rock et al., 2017). Finally, we predicted that Susceptible rats would demonstrate greater anxiety in the light-dark box than would Resilient rats.

## Methods

### Animals

Adult male Sprague-Dawley rats (Charles River; N = 307) were individually housed in ventilated cages in a vivarium maintained on a 12:12 light-dark cycle (lights off at 7:00 pm). Prior to the beginning of the study, rats were acclimated to the vivarium for 7 days with ad libitum access to food and water. Beginning 72 hours after arrival, rats were carefully handled to become familiar with experimenters (always one male and one female experimenter) prior to stress induction. Food access was restricted to 20 g/day from the beginning of testing until the end of experiments. All procedures were performed within 4 hours of the beginning of the dark cycle. Rats arrived in 3 cohorts of between 80-110 rats over the course of one year. Procedures were approved by the Institutional Animal Care and Use Committee at the University of Florida.

### Drugs

3-Cyano-N-(1,3-diphenyl-1H-pyrazol-5-yl)benzamide (CDPPB, 30 mg/kg; Abcam Biochemical) was suspended in 10% Tween 80 (Sigma-Aldrich) in phosphate-buffered saline (PBS) to a final concentration of 30 mg/ml and injected subcutaneously (s.c.). The dose of CDPPB was based on previous studies indicating an effect on fear extinction (Sethna & Wang, 2014). Cannabidiol (CBD, 5 mg/kg) was provided by the NIDA controlled substances program (RTI, Research Triangle, NC) and dissolved in a mixture of 100% ethanol, Cremophor, and 0.9% NaCl to 5 mg/ml and injected intraperitoneally (i.p.). The dose of CBD was based on previous studies demonstrating an effect on enhancing footshock conditioned contextual fear conditioning (Jurkus et al., 2016), and is within a range of doses found to produce anxiolytic effects (Guimarães et al., 1990). Locomotor testing was not performed as given doses of CDPPB and CBD do not affect locomotion in rats (Gass & Olive, 2010; Ren et al., 2010). 2, 3, 5-Trimethyl-3-thiazoline (TMT, 5 μl; BioSRQ) was presented undiluted (97% purity). The amount of TMT used for predator odor exposures was based on previous studies by our laboratory and others (Day et al., 2004; Tanapat et al., 2001; Schwendt et al., 2018).

### Experimental Procedures

#### Predator-scent stress induction and assessment of anxiety

The timeline for the predator-scent stress exposure and assessment of anxiety is shown on Fig 1A. Six to ten days after arriving in the vivarium, rats received a single exposure to the fox odor TMT. These exposures occurred in a covered, clear cylindrical Plexiglas chamber (BioBubble Pets; 40 cm diameter x 35 cm height) where the steel mesh flooring was raised to allow for a clear plastic dish to be placed underneath. Prior to each session, fresh TMT (5 μl) was pipetted onto a small square of filter paper which was positioned in the center of the dish. Rats were individually placed in the test chamber for a single 10 min exposure. Test chambers were cleaned with 70% ethanol between sessions.

**Figure 1.**
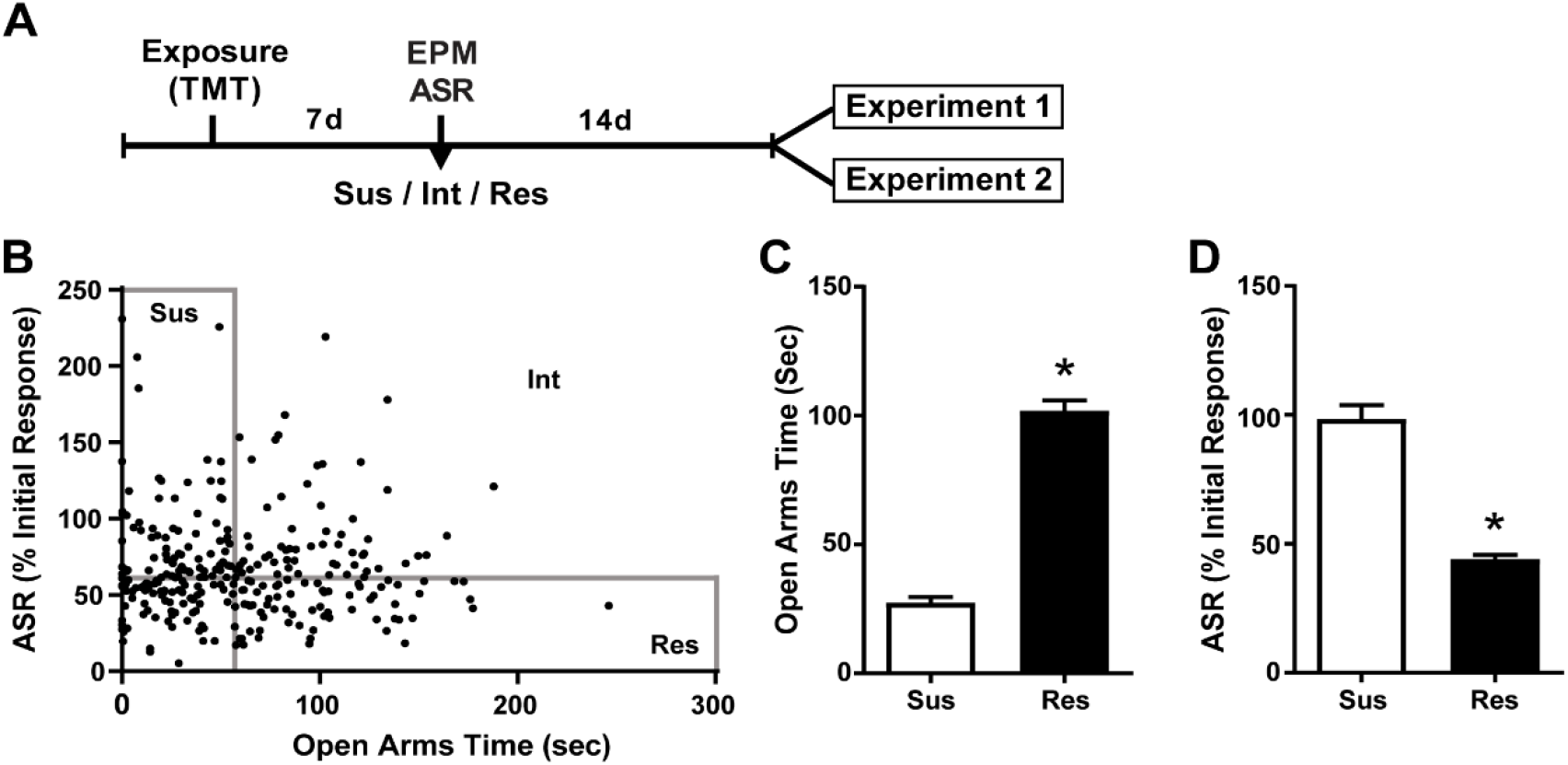
Susceptible and Resilient rats display distinct behavioral phenotypes. **A**, Timeline for behavioral classification. **B**, Time spent in the open arms of the EPM plotted against % habituation of acoustic startle response (ASR) for all TMT exposed rats (*n* = 299). Median splits performed on EPM open arm time (median = 56.2 sec) and % ASR habituation (median = 61.2%) were used to classify rats into Susceptible (Sus, *n* = 74), Resilient (Res, *n* = 74), or intermediate (Int, *n* = 151) phenotypes. **C**, Sus rats spent less time in the open arms of the EPM and d, exhibited attenuated habituation to ASR relative to Res rats. \**p* < 0.0001.

Seven days after TMT exposure, rats were tested on the EPM according to previously described procedures (Pellow et al., 1985). The EPM apparatus (Med Associates) was made from black acrylic and consists of four arms (50 cm length x 10 cm width) raised 50 cm from the floor. Two open arms (2.5 cm high walls) and two closed arms (50 cm high walls) are joined by a center square platform (10 cm x 10 cm) illuminated at 50 lux. Rats were individually placed on the center platform facing a closed arm and allowed to move freely for 5 min. Sessions were filmed by a camera secured above the maze. The EPM was cleaned with 70% ethanol between tests. Total time spent in the open arms excluding time in the center area (OA time) was recorded with EthoVision XT 14 software (Noldus Information Technology) and served as a measure for anxiety.

Immediately after EPM testing, habituation of acoustic startle response (ASR) was assessed according to Valsamis & Schmid (2011). Four ventilated soundproof chambers (San Diego Instruments) each contained a transparent plexiglass cylinder that rested on a pressure-sensitive platform. An accelerometer fitted to the platform measured changes in pressure created from movement of the rat’s body, and the maximum response amplitude was registered during presentation of acoustic stimuli. Accelerometer calibration and acoustic sound levels were routinely checked, and chambers were cleaned with CaviCide disinfectant (Metrex) and 70% ethanol between sessions. Rats (four at a time) were secured in the plexiglass cylinders and acclimated to the chamber for 5 min. Next, 30 pulses of 110 db white noise were delivered for 40 ms followed by a variable (30-45 s) intertrial interval. Startle habituation was calculated as the percent change in startle amplitude from the first six trials to the last six trials.

Power analyses (G power) indicated the number of animals needed was 7-8/ treatment group with a significance level of 0.05. We have previously shown that the incidence of the Susceptible phenotype amongst TMT-exposed rats ranges from 14-21.8% (Schwendt et al., 2018). Thus, with a target of 8 rats/group for a total of 16 Susceptible rats needed for Experiment 1 and 32 needed for Experiment 2, we initially exposed 307 rats to TMT.

#### Experiment 1 – The effects of CDPPB of contextual fear extinction and context-induced Fos protein expression

The timeline for this experiment is shown in Figure 2A. Rats first underwent predator stress induction and anxiety assessment (see above and Fig. 1A). Rats classified as Susceptible were randomly subdivided into two groups: Sus-Veh and Sus-CDPP (*n* = 7/group). Two weeks after anxiety assessment and phenotype classification rats underwent three contextual fear extinction sessions once daily. We previously found that in Susceptible rats with a history of cocaine self-administration, treatment with CDPPB immediately prior to placement into the TMT context increased freezing on days 2-4 of extinction (Schwendt et al., 2018). Here we sacrificed rats after only three days of extinction in order to examine neuronal activity during the last extinction session. Twenty minutes before sessions, rats were injected in their home cage with either vehicle or CDPPB. Rats were then placed into the exposure chamber in the absence of TMT for 10-minute sessions (one session/day). To avoid potential residual scent, we used plexiglass chambers which never had contact with TMT. Sessions were filmed, and freezing was quantified offline using the mobility detection function in Ethovision XT 14 software according to Pham et al. (2008). Freezing in rats is a species-specific threat-related defensive strategy that is defined by the absence of movement except for respiration (Fanselow, 1980).

**Figure 2.**
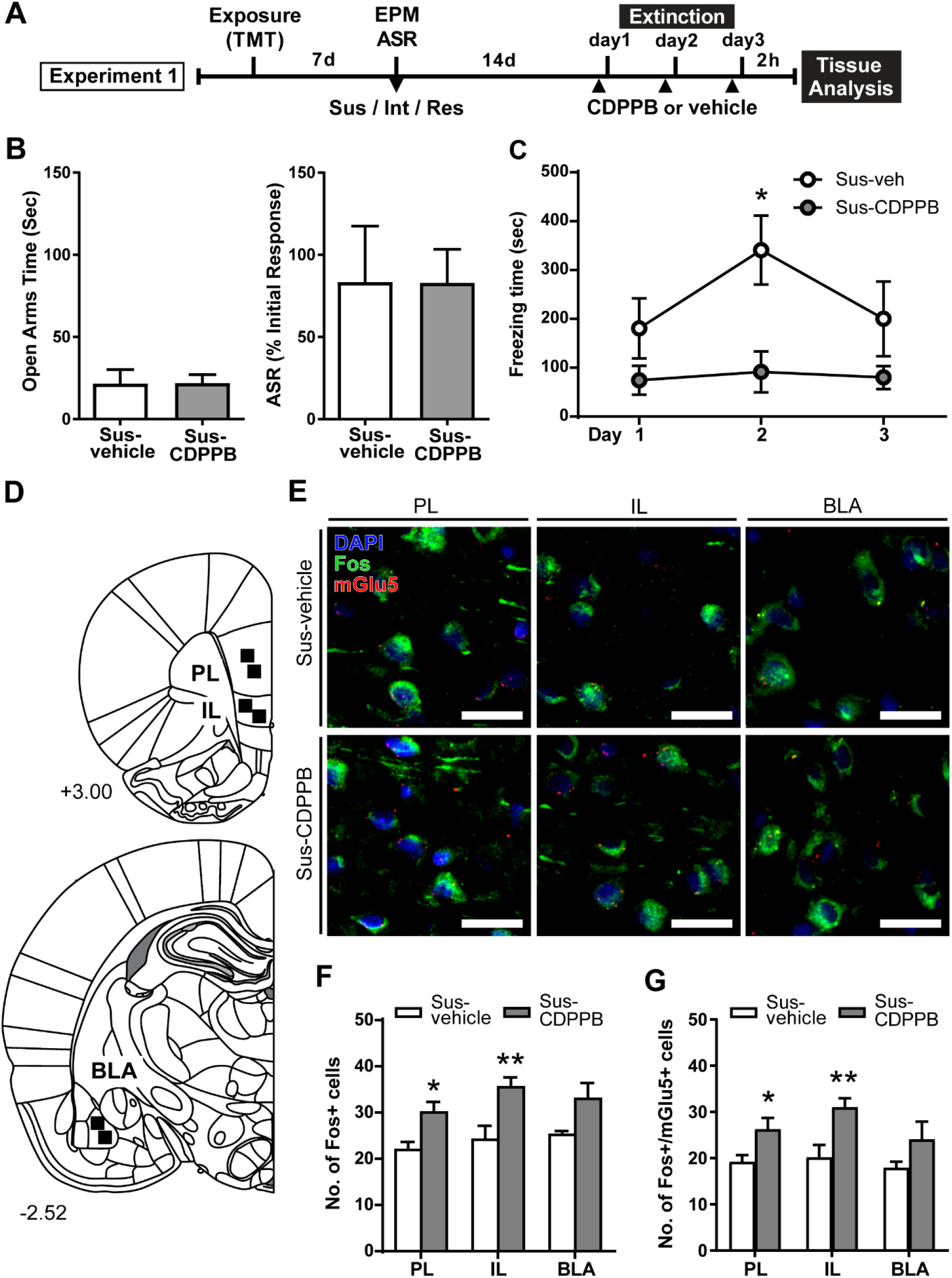
CDPPB reduces context dependent freezing in Sus rats and increases Fos expression in the medial prefrontal cortex. **A**, Timeline for Experiment 1. **B**, Treatment groups did not differ in EPM open arm time or % ASR habituation. **C**, Mean ± SEM of freezing in vehicle (*n* = 7) or CDPPB (*n* = 7) treated Susc rats during 10 min extinction sessions in the TMT context initiated 14 days after phenotype classification; CDPPB reduced freezing on day 2 of extinction. **D**, Schematic indicating the regions of interest (ROI) used for characterization of extinction dependent neuronal activity in the prefrontal cortex (top) and basal lateral amygdala (BLA; bottom). **E**, Representative confocal images from the selected ROIs in the prelimbic cortex (PL; left), infralimbic cortex (IL; middle) and BLA (right) showing fluorescent immunolabeling of Fos (green) protein, labeling of mRNA for mGlu5 using fluorescent in situ hybridization, and DAPI (blue) nuclear staining, in vehicle (top panel) and CDPPB (bottom panel) treated susceptible rats; scale bar: 20 μm, 63x magnification. **F**, CDPPB increased the number of Fos labeled cells in the PL and IL regions on day 3 of extinction. **G**, The effects of CDPPB on the number of Fos+/mGlu5+ double-labeled cells in the PL, IL and BLA on day 3 of extinction. \**p* < 0.05, \*\**p* < 0.01 relative to vehicle.

Next, we performed fluorescent immunolabeling for Fos protein to measure neuronal activity during the third extinction session. Fos is widely-used relative marker for neuronal activation (Kovacs, 2008; Lanahan & Worley, 1998). As previous studies have demonstrated a role for mGlu5 in the regulation of cFos, and stimulation of mGlu5 with CDPPB can enhance cFos expression (Mao et al., 2005, 2014; Uslaner et al., 2009), we also performed dual-labeling for mGlu5 mRNA. Two hours after the third extinction session, rats were administered an overdose of sodium pentobarbital (Euthasol, 1 ml, i.p.) and transcardially perfused in nuclease free 0.9% NaCl followed by cold 4% paraformaldehyde (PFA) in PBS. Brains were removed, post-fixed for 12 hrs at 4°C in 4% PFA in PBS, cryopreserved in 30% sucrose in PBS, frozen and kept at −80°C until sectioning. Twelve-μm-thick tissue sections corresponding to the rat PL/IL (+3.00 mm relative to Bregma) and BLA (−2.52 mm) according to rat brain atlas (Paxinos & Watson, 2005), were collected using a freezing cryostat (Leica CM1950). Tissue sections were direct-mounted onto Superfrost Plus Gold slides (Fisher Scientific), dried and stored at −80°C. Fluorescent in situ hybridization of *GRM5* (mGlu5) mRNA was performed using the RNAscope Multiplex Fluorescent Reagent Kit (ACDBio) according to the manufacturer’s instructions with a few modifications (Wang et al., 2012). Slides were first equilibrated to room temperature (RT) before heating to 60°C for 45 min. Next, sections were fixed in cold 4% PFA in PBS and dehydrated using ethanol gradient of 50%, 70%, 100%, 100% in consecutive 5 min incubations. Slides were boiled in target retrieval reagent (ACDbio), washed in nuclease-free H2O, and again dehydrated in 100% ethanol. Proteinase digestion of sections was conducted using pretreatment #3 (ACDbio) at 40°C for 30 min under humidity-controlled conditions (HybEZ hybridization owen; ACDbio). The RNAscope probe for *GRM5* (ACDBio: 471241-C2, lot 17335A) was diluted with C1 diluent probe (1:50) and applied to sections. Slides were then incubated 2 h at 40°C for hybridization of probe to target mRNAs. Signal amplification was performed with preamplifier and amplifier probes at 40°C (AMP 1, 30 min; AMP 2, 15 min; AMP 3, 30 min; AMP 4, 15 min). For AMP4 (15 min), Alt-A was selected so that the target probe could be detected with ATTO 550 (Cy3) fluorescent label. Immediately following in situ hybridization, slides were rinsed three times in Tris-buffered saline (TBS) and blocked (0.3% Triton x-100 and 5% NGS in TBS) for 1 hour at RT. TBS was used for all rinses and antibody dilutions. Sections were incubated in rabbit anti-Fos antibody (1:1000; Synaptic Systems) overnight at 4°C. The next day slides were rinsed and then incubated in goat anti-rabbit Alexa 594 secondary antibody (1:1000; Invitrogen) for 2 h at RT. Finally, sections were rinsed again before counterstaining with DAPI (Invitrogen) and coverslipped using ProLong Gold antifade mounting reagent (Invitrogen). Fluorescent images were captured using Zeiss LSM70 inverted Axio-Observer 3-channel spectral confocal microscope and Zen 2012 software. Multitrack sequential imaging settings were applied to avoid inter-channel crosstalk effects. The 405, 488, and 561 nm laser lines were used for excitation of DAPI, c-Fos (Cy3), and mGlu5 (FITC). Two regions of interest were selected for each brain area (PL, IL, and BLA; Fig. 2D.) and Z-stacks were acquired at 1μm intervals using a 63X oil immersion objective. Image stacks were imported into NIH Image J software (Schneider et al., 2012) and analyzed offline. Only the middle five focal planes from each Z-stack were used for analysis. To measure co-expression, Z-stacks were first converted to composite images for separation of individual color channels. The red channel (mGlu5) was then turned off, and cells containing green (Fos) staining were marked using the NIH Image J multi-point tool on each focal plane. Then, the green channel was then turned off, and the red channel was used to mark mGlu5 puncta in the same manner. Individual cells were distinguished based on DAPI nuclei staining. Fos expression was established if staining within a cell was detectable on three consecutive focal planes, and mGlu5 expression was established if a cell contained mRNA puncta on three consecutive focal planes. Under these conditions, cells expressing both Fos and mGlu5 were considered positive for co-expression. The total number of Fos expressing and Fos/mGlu5 co-expressing cells were counted for each selected area. The average number of cells for each parameter across the two selected areas was calculated for the three brain regions for each rat.

#### Experiment 2 – The effects of CDPPB (or CBD) on unconditioned anxiety and contextual fear extinction

The timeline for this experiment is shown in Figures 3A and 4A. Rats first underwent predator stress induction and anxiety assessment as described above. Susceptible rats were randomly selected from the sample to create vehicle (Sus-Veh; *n* = 8), CDPPB treated (Sus-CDPPB; *n* = 8), and CBD treated (Sus-CBD; *n* = 8) groups. Resilient rats were selected to create a vehicle treated (Res-Veh; *n* = 8) group. Two weeks after anxiety assessment and phenotype classification, rats underwent light-dark box testing based on previously described procedures (Crawley, 1981). Light-dark box apparatus consisted of a plexiglass box with two separate compartments of equal dimensions (40 cm length × 44 cm width x 37 cm; a light compartment with translucent walls illuminated at 300 lux, and a dark compartment with blackened opaque walls. The compartments were connected by an opening in the dividing wall, allowing rats a free transition between the boxes. Prior to testing, rats were treated with vehicle, CDPPB, or CBD in their home cage. Following pretreatment (CDPPB: 20 min; CBD: 30 min), each animal was individually placed in the center of the light box and allowed to roam freely for 10 min. Sessions were filmed, and the following behaviors were hand scored by an experimenter blind to the conditions: (1) latency to enter the dark box from the light box and vice versa; (2) the no. of transitions from the dark box to the light box; and (3) the duration of individual compartment visits. The apparatus was cleaned with 70% ethanol between trials. Next rats were used to assess differences between Resilient and Susceptible rats in fear extinction and the ability of CBD to enhance the extinction of TMT conditioned fear. The day after light-dark box testing, rats were administered vehicle or CBD 20 min prior to testing (Fig. 4A). Rats received the identical treatment that they had received on the day prior for light-dark box testing. Procedures for fear extinction were identical to Experiment 1.

**Figure 3.**
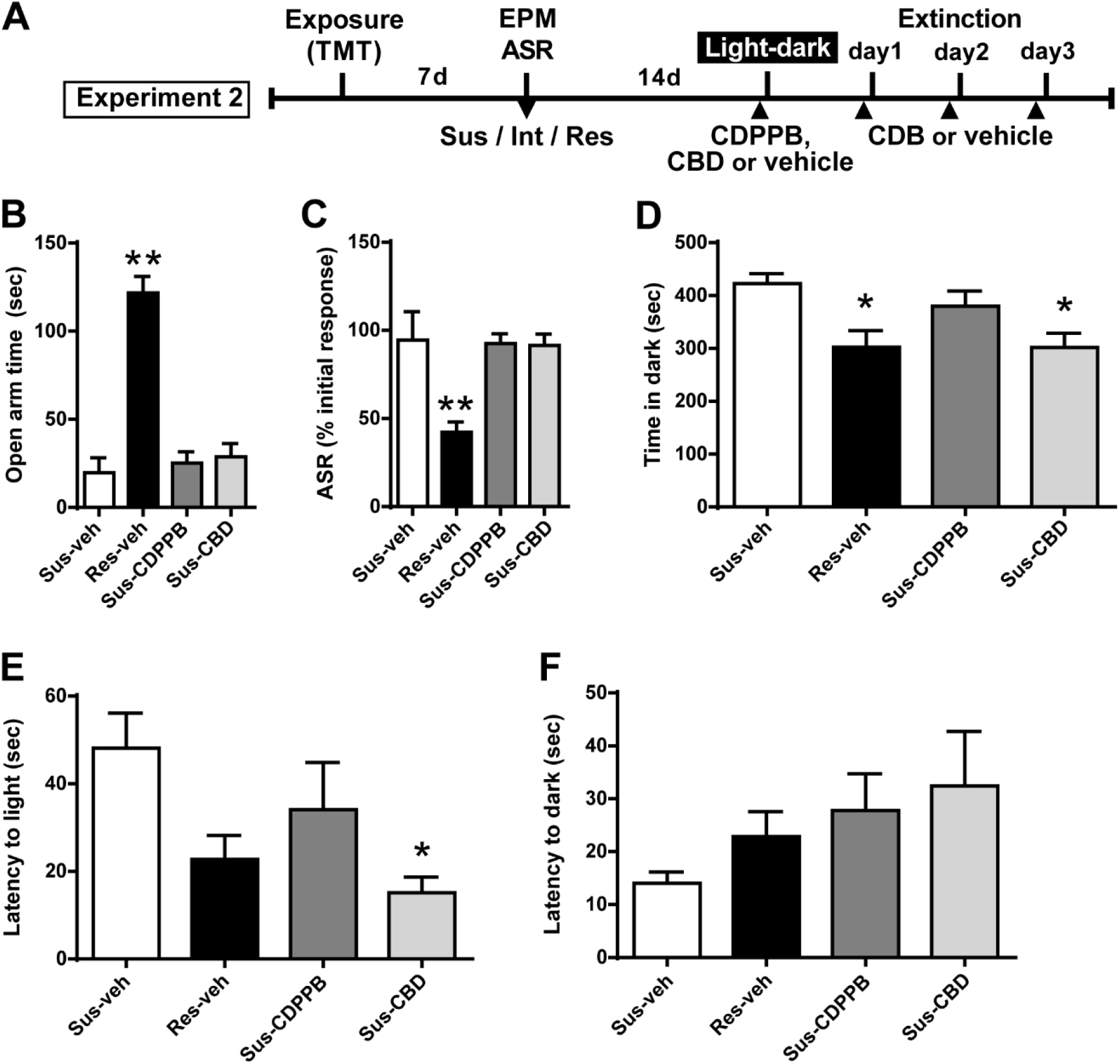
CBD attenuates increased anxiety-like behavior in the light-dark box test in Susceptible rats. **A**, Timeline for Experiment 2. **B**, Vehicle (Sus-veh, *n* = 8), CDPPB (Sus-CDPPB), and CBD (Sus-CBD, *n* = 8) treated Sus rats spent less time in the open arms of the EPM and **C**, exhibited reduced habituation to ASR relative to vehicle treated Res rats (Res-veh, *n* = 8). **D**, CBD treated Sus rats and Res rats spent less time in the dark-box relative to vehicle treated Sus rats. **E**, CBD treatment reduced the latency to enter the light-box in Sus rats. **F**, No differences were observed in latency to enter the dark-box. \*\**p* < 0.05 relative to Res-veh; *p < 0.05 relative to Sus-veh.

**Figure 4.**
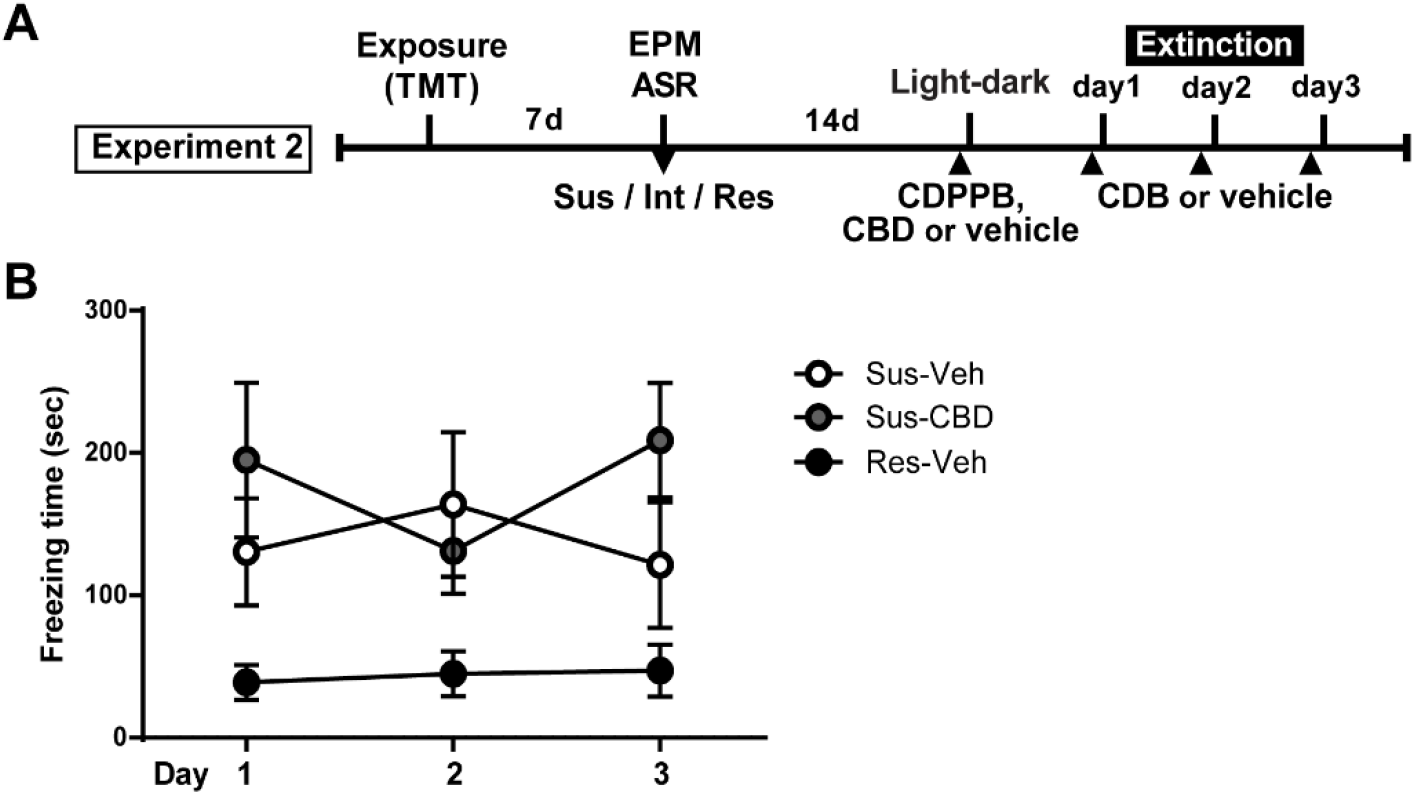
CBD does not reduce increased context dependent freezing in Susceptible rats. **A**, Timeline for Experiment 3. **B,** Mean ± SEM of freezing in vehicle (Sus-veh; *n* = 8) and CBD (Sus-CBD; *n* = 8) treated Sus rats and vehicle treated Res rats (Res-veh; *n* = 7) during 10 min extinction sessions. Sus-veh and Sus-CBD displayed increased freezing relative to Res-veh rats. *main effect of phenotype

### Data analysis

GraphPad Prism (version 6.0) was used for statistical analysis with the alpha level set at p ≤ 0.05. Unpaired t-tests were used to compare phenotypic differences in EPM and ASR behavior for rats used in Experiment 1-2 and for within region comparison of neuronal markers between treatment groups. One-way analyses of variance (ANOVAs) were used to measure phenotypic differences in EPM and ASR behavior for rats used in Experiments 3-4, and to test for phenotype/treatment effects on light-dark box testing. Fear extinction was analyzed by two-way mixed factorial repeated-measures (RM) ANOVAs with Treatment as the between-subjects factor and Day as a within-subjects factor. Significant interactions were followed by Tukey’s post hoc analyses with corrections for multiple comparisons.

## Results

### Predator odor stress and susceptibility classification

A total of 307 rats were exposed to TMT. ASR data files for 8 rats were corrupted and thus the following calculations are based on the data from 299 rats. The median time spent in the OA of the EPM was 56.2 seconds and median habituation of the ASR was 61.2%. Rats that fell below the median for time spent in the OA and above the median for ASR habituation were classified as Susceptible as in our previous work (Schwendt et al., 2018).

Rats were classified as Resilient if they fell above the median for time spent in the OA and below the median for ASR habituation. This resulted in 74 rats (25%) meeting criteria for the Susceptible (Sus) phenotype and an equal number meeting criteria for the Resilient (Res) phenotype (Fig. 1B). A total of 46 Susceptible rats and 8 Resilient rats were used for the present set of experiments. More Susceptible and Resilient rats were generated than were needed for the present set of experiments and were used for other experiments to be published at a later date. Rats not classified as either Susceptible or Resilient (Intermediate; *n* = 151) were eliminated from the experiment.

### Experiment 1

Susceptible (Sus) rats spent less time in the open arms of the EPM [*t*(29) *t*(26) = 4.482 p < 0.0001, Fig. 1C], and exhibited less reductions in ASR magnitude [*t*(26) = 4.515, p < 0.0001, Fig. 1D] as compared to Resilient rats. The two treatment groups of Susceptible rats (Sus-vehicle, *n* = 7; Sus-CDPPB *n* = 7) used for Experiment 1 did not significantly differ in EPM and ASR scores (Fig. 2B,C). A 2-way RM ANOVA conducted on freezing behavior (Fig. 2A) revealed a main effect of Treatment [*F*(1, 11) = 6.803, *p* = 0.024] and Day [*F*(2, 22) = 3.905, *p* = 0.035]], and a significant Treatment x Day interaction [*F*(2, 22) = 5.134, *p* = 0.015]. Post hoc analysis revealed that on day 2, Sus-CDPPB rats froze significantly less than Sus-veh rats, and no differences in behavior were observed on days 1 or 3 (Fig. 2C). Analysis of Fos immunoreactive cells revealed increased Fos expression in the PL [t(10) = 2.80, p = 0.02], IL [t(10) = 3.03, p = 0.01] of Sus-CDPPB rats compared to Sus-veh (Fig. 2F). In the BLA, a trend towards increased Fos protein expression was detected in Sus-CDPPB group (p = 0.054). Co-expression of Fos and mGlu5 was abundant across analyzed regions of both treatment groups. However, the level of Fos/mGlu5 expression overlap varied between regions (Sus-veh, [*F*(2, 15) = 2.756, *p* = 0.0125]; Sus-CDPPB [*F*(2, 15) = 0.830, *p* < 0.0001]), with lower co-expression in the BLA (∼70%) compared to both IL and PL regions (∼90%; data not shown). Perhaps due to this high degree of co-localization, CDPPB treatment increased Fos expression in mGlu5-positive cells in a manner that was similar to overall Fos expression: expression was increased in the PL [t(10) = 2.92, p = 0.04] and IL [t(10) = 3.30, p = 0.01], nut not in the BLA (Fig. 2G). Representative images of dual Fos/mGlu5 labeling from each of these regions are shown in Figure 2E.

### Experiment 2

Subsets of Susceptible (Sus) and Resilient (Res) rats were used to generate four treatment groups (Sus-veh, Sus-CDPPB, Sus-CBD, Res-vehicle). A one-way ANOVA revealed phenotypic differences in EPM open arm time [*F*(3, 27) = 37.61, *p* < 0.0001, Fig. 3B] and ASR habituation [*F*(3, 27) = 8.028, *p* = 0.0006]. All three groups of Sus rats spent less time compared to Res rats in the open arms of the EPM (Sus-veh, *p* < 0.0001; Sus-CDPPB, *p* < 0.0001; Sus-CBD, *p* < 0.0001, Fig. 3B), and exhibited less habituation of the ASR relative to Res rats (Sus-veh, *p* = 0.0023; Sus-CDPPB, *p* = 0.0023; Sus-CBD, *p* = 0.0029, Fig. 3C). In the light-dark box test, Group differences in time spent in the dark side were found [*F*(3, 27) = 4.686, *p* = 0.0092, Fig. 3D]. Post hoc analysis revealed Sus-veh rats spent more time in the dark side compared to Res-veh, indicating the preservation of anxiety phenotypes two weeks after classification with EPM and ASR. In Susceptible rats, CDPPB did not alter time spent in the dark (*p* = 0.692, Sus-CDPPB vs. Sus-veh), however CBD had an anxiolytic effect, demonstrated by a reduction in dark box time (*p* = 0.023, Sus-CDB vs. Sus-veh). Time spent in the dark in Sus-CBD rats did not differ from Res-veh rats. One-way ANOVA also revealed differences in latency to enter the light box [*F*(3, 25) = 3.315, *p* = 0.036, Fig 3E], with CDPPB again having no effect, and CBD reducing the time to enter relative to the Sus-veh treated group. No phenotypic or treatment differences were observed in the latency to enter the dark side. A two-way repeated measures ANOVA on time spent freezing revealed a significant main effect of group [*F (2, 20) = 4.106, p* = 0.032] and a trend towards a significant group x time interaction [*F (4, 40) = 2.197, p* = 0.08]. There was no main effect of time [F (2, 40) = 0.2389, n.s.] (Fig. 4B).

## Discussion

Here we demonstrate that seven days after a single exposure to the predator odor TMT, populations of stress Susceptible and Resilient rats emerged following phenotype classification using EPM and ASR scores. These findings conform to previous reports demonstrating lasting manifestations of anxiety in subpopulations of predator odor-exposed rodents (Cohen & Zohar, 2004; Schwendt et al., 2018), and correspond with heightened anxiety and startle-response observed in human PTSD (Shalev et al., 1992). As in our previous report (Schwendt et al., 2018), we classified rats with a double median split that required an anxiogenic response across both tasks to meet criteria for the Susceptible phenotype and the opposite to be classified as Resilient. The majority of rats displayed EPM and ASR behavior that was intermediate between Susceptible and Resilient rats, signifying that the two phenotypes were indeed representative of extremes on the susceptibility spectrum. In addition, this study revealed that anxiety behavior in the light-dark box is consistent with prior phenotype characterization, such that Resilient rats displayed less anxiogenic behavior than Susceptible rats, indicating that both phenotypes are stable for at least three weeks after TMT exposure. This supports our previous observations that rats meeting the criteria for the Susceptible and Resilient phenotypes exhibit distinct differences in cortisol concentrations, freezing in the TMT context, gene expression, and cocaine seeking that persist for two to five weeks after EPM and ASR classification (Schwendt et al., 2018).

In agreement with our previous report (Schwendt et al., 2018), in the current study we found a conditioned fear response in Susceptible rats three weeks following the initial TMT context pairing that was greater than that in Resilient rats, and this conditioned fear response did not extinguish over the course of three days. This is consistent with studies that demonstrated the presence of robust defensive behaviors in rodents following re-introduction to a TMT paired context (Endres & Fendt, 2007; Rosen et al., 2008). While some previous studies failed to produce conditioned fear response with TMT (McGregor et al., 2002; Blanchard et al., 2003; Rosen, 2004), these studies only tested for such a response 24 hours after the exposure. It has been proposed that TMT may have a weak capacity to serve as a predictive cue for a predator, and paradigms successful in producing TMT conditioned fear response focused on manipulating the exposure apparatus to increase its ability to facilitate salient TMT-context associations (Takahashi et al., 2008; Rosen et al., 2015). We hypothesize that conditioned fear responses in the TMT-associated context were reported here and in our previous publication due to the inclusion of a post-stress incubation period, and longer exposure time in a medium size chamber which facilitated the accumulation of sufficient contextual CS information and allowed for display of appropriate defensive behavior (freezing). Furthermore, our group is the first to segregate stress-susceptible and resilient rats, and susceptible rats are likely more vulnerable to conditioning. It is important to note that none of the Susceptible rats extinguished freezing here. Thus, our model captures both the lasting manifestations of anxiety and extinction deficits observed in individuals with PTSD.

Consistent with previous rodent studies using conditioned footshock (Sethna & Wang; 2014), we found that the mGlu5 PAM CDPPB facilitated extinction in Susceptible rats, suggesting common mechanisms are involved in extinction of innate predator fears and discrete aversive stimuli. These findings support growing evidence implicating mGlu5 in the modulation of fear memory (Xu et al., 2009; Fontanez-Nuin et al., 2011). Consistent with our previous report, we observed an increase in freezing in vehicle-treated rats from Day 1 of extinction to Day 2, indicating an incubation of fear (Schwendt et al., 2018). In agreement with an established role for CDPPB in the enhancement of memory consolidation (Sethna & Wang, 2014; Uslaner et al., 2009), CDPPB treated rats demonstrated less freezing on the second day of extinction compared to vehicle rats. It is interesting to note that rats treated with CDPPB showed similar freezing behavior as did Resilient animals, which as reported in our previous study, show an enhanced mGlu5 mRNA expression in the mPFC and amygdala. These findings further suggest a role for mGlu5 function in resilience to predator odor trauma. However, while we previously observed that CDPPB increased freezing during days two and three of fear extinction in rats with a history of cocaine self-administration, here we found that CDPPB treated rats froze less than vehicle rats on day 2, indicating that chronic self-administration of cocaine alters the neurobiology underlying the extinction of conditioned fear. Interestingly, previous studies in rats and humans have demonstrated bi-phasic effects of the NMDA receptor partial agonist D-cycloserine, which can either produce a weakening or a strengthening of fear memory contingent upon an individual’s experience during the extinction session (Orr et al., 2000; Bolkan & Lattal, 2014). Furthermore, in cocaine users, D-cycloserine prevents reductions in craving and brain activation produced by extinction of cocaine-associated cues. Taken together, this suggests that cocaine alters the role of glutamate receptors in mediating extinction of both fear and cocaine-associated cues (Prisciandaro et al., 2013; Price et al., 2012). This should be considered for the treatment of anxiety disorders in patients with cocaine use disorder.

Although we did not observe differences in freezing between treatment groups on extinction day 3, Fos immunolabeling revealed increased IL and PL activity in CDPPB-treated rats. Analysis of Fos+mGlu5 co-expression revealed that the increased Fos expression was most likely a result of CDPPB, as a) very few Fos-positive cells did not contain mGlu5 mRNA, and b) increase in Fos expression in mGlu5-positive cells mirrored overall Fos protein increase. This increase is likely a result of extinction-mediated Fos activation in combination with direct pharmacological effects of CDPPB, as activation of mGlu5 receptor signaling increases the phosphorylation of CREB, a transcription factor for the *c-Fos* gene (Uslaner et al., 2009, Wang & Zhuo, 2012). In relation to this, a lack of CDPPB effect, in the BLA can be attributed to a ‘ceiling effect’, as animals with fear-extinction deficits display a robust Fos expression in the BLA (Hoffman et al. 2012).

That IL activity was enhanced in our CDPPB treated rats is consistent with the growing body of work implicating a central role for the IL in fear extinction (Do-Monte et al., 2015; Herry et al., 2008; Cho et al., 2013). During extinction, glutamate driven neuronal activity within the IL triggers synaptic enhancements across IL–amygdala inhibitory circuits, effectively suppressing CeA output cells and preventing behavioral fear response. Thus, it is possible that if in the CDPPB treated rats a similar rise in IL activity as the one seen on day 3 occurred during day 2, it might have contributed to the inhibition of fear as compared to vehicle treated rats. However, this does not support the absence of a difference in fear response on day 3 regardless of increased IL activity. This discrepancy might be explained by our observation that both IL and PL activity were increased during the session. Although PL activity has been observed during extinction of footshock conditioned cues (Wilber et al., 2011; Knapska & Maren, 2009), a reduction in PL activation is more commonly demonstrated, as PL signaling is associated with extinction resistance (Burgos-Robles, 2009). It may be the case that although CDPPB enhanced neuronal activity in the IL, effectively promoting the inhibition of fear response, these benefits were countered by the nonselective targeting of the PL region. It is important to note that evidence for opposing roles of the IL and PL during extinction has been exclusively characterized in footshock models. Interestingly, the only other study measuring neuronal activity following predator odor extinction observed similar increases across the entire mPFC, suggesting the possible involvement of an alternative circuitry (Mackenzie et al., 2010).

While mGlu5 may be a promising target for extinction enhancements, it is important to consider previous studies demonstrating that mGlu5 agonists can produce anxiogenic effects, and antagonists are anxiolytic (Rahman et al., 2017; Klodzinska et al., 2004). Here, we found that CDPPB neither attenuated nor enhanced anxiety in the light-dark box task. This is in contrast to a recent report that the same dose of CDPPB increased anxiety measures in the light-dark box in unstressed mice and in mice that had consumed alcohol (Lee et al., 2018). Thus, CDDPP or other mGlu5 PAMs may be beneficial in a PTSD population as they would enhance fear extinction without inducing general anxiety, but caution should be exerted in alcohol users.

Although prior studies in heterogeneous rodent populations have observed extinction enhancing properties of CBD in a conditioned footshock model (Stern et al., 2012; Bitencourt et al., 2008; Do-Monte et al., 2013), we did not observe an effect across five sessions of extinction in our susceptible rats. It is possible that TMT exposure was not sufficient in evoking the neural adaptations which benefit from CBD mediated singling. Indeed, one recent study demonstrated that while CBD enhanced extinction in footshock conditioned rats, extinction was unaffected by CBD in rats administered a reduced number of shocks (Song et al., 2014). It may also be the case that our dose of CBD was not strong enough to effectively enhance or impair fear extinction, as 5 mg/kg is on the lower end of the inverted U dose response curve for extinction enhancing effects (Song et al., 2014; Zuardi et al., 2017). The mechanisms by which CBD exerts pro-extinction and anti-anxiety effects are multifaceted, however several studies have demonstrated a capacity for direct and/or indirect potentiation of CB1R (Bisogno et al., 2001; Campos et al., 2013). Endocannabinoid signaling (eCB) is profoundly implicated in fear extinction through its involvement in synaptic plasticity (Heifets & Castillo, 2009), and dysregulated eCB signaling has been implicated in PTSD (Neumeister et al., 2013). In addition, CBD can function as a partial agonist of the 5-HT1a receptor (Russo et al., 2005; Campos et al., 2012b; Fogaca et al., 2014)). Abnormal 5-HT signaling has long been understood to contribute to PTSD like symptoms (Zhao et al., 2017), and is a commonly used target for treatment of the disorder (Sullivan et al., 2013). Thus, further study is needed to more exhaustively characterize the potential of CBD in PTSD treatment. While CBD did not enhance extinction, it was effective in reducing anxiety in the light-dark box test. Our findings are consistent with previous work showing benefits of CBD in alleviating heightened anxiety in animals that have experienced prior footshock or restraint-stress (O’Brien et al., 2013; Song et al., 2014; Fogaça et al., 2014). While chronic CBD injections following exposure to a live predator was effective in mitigating predator-induced anxiety one week after exposure (Campos et al., 2012), we are the first to show an effect of CBD with only a single acute dose in a population of Susceptible rats. Our findings are consistent with previous studies demonstrating an anxiolytic effect of CBD only in animals previously exposed to stressors (Rock et al., 2012; O’brien, 2013 Restell et al., 2009; Bitencourt et al., 2008).

## Conclusions

Here we found strong evidence that mGlu5 PAMs, such as CDPPB, represent a potential treatment strategy for the enhancement of extinction of conditioned fear in a PTSD population. Such enhancement of extinction comes without increased anxiety, which has previously been reported for this class of drugs in some population of animals. Based on preclinical work (Schwendt et al., 2018; Lee et al, 2018), caution should be exerted in when using this class of drugs in patients including with a history of cocaine or alcohol use. Finally, while CBD was anxiolytic, it failed to enhance the extinction of conditioned fear in a Susceptible, PTSD-like population of rats. Future consideration should be given to polytherapy with both CBD and mGlu5 PAM for the enhancement of extinction and relief of anxiety that accompany PTSD.

## Acknowledgements

We would like to thank Lizhen Wu, Daniel Hopp and Stephen Beaudin-Curley for their assistance with behavioral experiments and/or analysis of behavioral data.

## Author contributions statement

JS, MS, & LK were involved in the conception and design of the study. JS, PH, MR and ARB were involved in the acquisition of data. JS, ARB, LK, and MS were involved in analysis and interpretation of the data. JS drafted the manuscript, and LK and MS made critical revisions. JS,PH, ARB, MR, LK & MS approved the final version of the manuscript to be published.

